# Pea-adapted biotype of the aphid *Acyrthosiphon pisum* induces susceptibility of pea to non-adapted biotype enabling improved feeding and performance

**DOI:** 10.1101/2022.09.12.507464

**Authors:** Po-Yuan Shih, Rémi Ollivier, Anas Cherqui, Arnaud Ameline, Stéphanie Morlière, Yannick Outreman, Jean-Christophe Simon, Akiko Sugio

## Abstract

Most insect herbivores are adapted to feed on a few host plants only, but the mechanisms underlying plant specialization are poorly understood. One of the dominant hypotheses is that insects inject an oral secretion into the plant that manipulates the plant’s defences resulting in induced susceptibility. We tested this hypothesis on the pea aphid (Acyrthosiphon pisum), which forms multiple biotypes, each specialized on specific legume hosts. In particular, we tested whether a pea-adapted biotype can induce host susceptibility on pea, which should positively affect the fitness of a non-pea-adapted biotype. We found that survival and fecundity of an alfalfa-adapted biotype on pea were enhanced by co-infestation but not pre-infestation with a pea-adapted biotype. The electrical penetration graph (EPG) method was then used to dissect the components of alfalfa biotype feeding behaviour on pea with and without co-infestation by the pea biotype. While probing time did not differ between these two conditions, differences were observed in most other EPG parameters. When co-infested with the pea-adapted biotype, the alfalfa-adapted biotype was more likely to establish a phloem feeding and exhibited a longer ingestion time on pea. Our results clearly indicated that the pea-adapted biotype induced pea susceptibility by increasing the accessibility of non-pea-adapted aphids to the host phloem sap, and as a result, the alfalfa biotype increased its survival rate and fecundity. These results contribute to understanding how aphids manipulate their specific host plants.

## Introduction

Plants are constantly challenged by pathogens and herbivores. Some herbivores simply consume stored nutrients, but others manipulate the plants and induce susceptible state to steal nutrients efficiently. Manipulation and induction of plant susceptibility are observed in some herbivorous insects and considered as one of the most aggressive strategies to consume plants (Karban & Agrawal, 2002). The manipulation may involve suppression of plant defence signalling or metabolomic changes, can be local or systemic, and is assumed to be induced by insect oral secretion or microbial symbionts (Chung et al., 2013; Karban & Agrawal, 2002; Musser et al., 2002; Robert et al., 2012; Sandström et al., 2000).

Aphids are sap-feeding insects. They insert specialised mouthparts, called stylets, to take phloem sap. During this process, aphids are known to secrete saliva that contains salivary effectors that alter the plant-aphid interactions (Pitino & Hogenhout, 2012; Shih et al., 2022). Induced susceptibility can be observed when aphids manipulate plant defences or nutrients through these effectors. A review of earlier works revealed that prior infestations of aphids generally increased the performance of aphids that were subsequently installed in the same leaves and in some cases, negatively impacted the performance of aphids installed in distant parts of the same plant (Züst & Agrawal, 2016). Local susceptibility can be induced also in the plants that have resistance genes against the aphids that attack them. For example, pre-infestation of a virulent biotype of the greenbug aphid, Schizaphis graminum, on resistant wheat increased the reproduction of subsequently installed avirulent biotype (Dorschner et al., 1987). Similarly, pre-infestation of a virulent clone of the peach-potato aphid, Myzus persicae, on pepper increased fecundity and survival rate of the avirulent aphids that were installed subsequently in the same site (Sun et al., 2020). Co-infestation experiments of the lettuce aphid, Nasonovia ribisnigri, on resistant lettuce showed that avirulent aphids installed in a group of virulent aphids increased their phloem feeding time and survival rate. In contrast, weak defence against virulent biotype was induced on both susceptible and resistant lettuce plants by avirulent biotype, which resulted in reduced phloem feeding time of the virulent biotype (Ten Broeke et al., 2017). In case of the soybean aphid, Aphis glycines, on resistant soybean, systemically induced susceptibility was observed upon aphid infestation by both virulent and avirulent biotypes: preceding infestation by either virulent or avirulent aphids increased fecundity of both virulent and avirulent aphids that were installed subsequently, and this result was not affected by aphid density (Varenhorst et al., 2015). These examples of induced-susceptibility show that each plant-aphid interaction involves specific defence and counter-defence mechanisms, and that it is difficult to generalize the results. Successive infestation tests may not be appropriate for revealing induced-susceptibility in some cases, as aphid-induced changes in plant biology may be transient and not last long enough to affect life-history traits that are classically measured (e.g. aphid survival or fecundity). Therefore, we assume that simultaneous infestation of two groups of aphids is more likely to detect induced effects, although it can be difficult to distinguish the two types of aphids.

Pea, *Pisum sativum*, is an important crop as it fixes atmospheric nitrogen to soil and provides high amount of proteins for human and animal consumption. However, it is constantly attacked by various parasites, such as the pea aphid, Acyrthosiphon pisum. The pea aphid forms a complex of at least 15 biotypes (Peccoud et al., 2015; Peccoud et al., 2009) and each biotype is specialised to feed on one or a few related legume species and cannot feed well on the others (Peccoud et al., 2009). For example, only the pea-adapted biotype (hereafter, pea biotype) can infest pea plants, whereas other biotypes, such as alfalfa-adapted biotype (hereafter, alfalfa biotype) cannot feed and develop on pea plants. The mechanisms that prevent the non-pea-adapted biotypes to feed on pea are not yet identified. An aphid behaviour study using the electrical penetration graph (EPG) showed that the pea biotype reached the phloem quickly and fed a long period, while the alfalfa biotype tried to feed on pea plant by inserting its stylets and salivating but could not establish phloem sap feeding (González et al., 2022; Schwarzkopf et al., 2013). We hypothesized that the differences in the feeding behaviour might be due to the biotype specific salivary proteins. In our previous work, 255 differentially expressed salivary genes were identified between *A. pisum* pea and alfalfa biotypes (Boulain et al., 2019). Some of those salivary proteins may supress the defence mechanisms of the specific host plants, such as production of toxic compounds to aphids and/or physiological changes (Shih et al., 2022). If so, the pea plants infested by pea-adapted aphids might induce local or systemic susceptibility that might be beneficial for the non-adapted biotypes. We also hypothesised the possibility that infestation of the alfalfa biotype on pea may trigger pea defence reactions and negatively affect the performance of the pea biotype.

Here, we first tested whether the pea biotype induces pea susceptibility and enhances the performance of the alfalfa biotype in the pre-infestation and co-infestation conditions. After observing positive effect of the co-infestation, we used EPG to record the feeding behaviour of the alfalfa biotype co-infested with the pea biotype to understand the mechanisms that improved its performance.

## Materials and MethodsPlants and aphids

Vicia faba (genotype Castel) and *P. sativum* (genotype AeD99OSW-50-2-5) (Ollivier et al., 2022) were used in this study, and both legume species were grown in a growth chamber set at 18°C with a 16 h day/8 h night photoperiod. The *P. sativum* genotype AeD99OSW-50-2-5 was chosen as it is highly resistant to the alfalfa biotype (Ollivier et al., 2022), which may increase the possibility of detecting the effect of induced susceptibility.

Three different clones of *A. pisum* were used including clone P123-amp of the pea biotype (green body colour), and clone LSR1 (pink body colour) and clone LL01 (green body colour) of the alfalfa biotype. All clones are deprived of secondary symbionts. LSR1 was the focal clone, used to detect pea susceptibility induced by clone P123. LL01, which is not expected to induce susceptibility in pea, was used to adjust aphid density in co-infestation experiment and to examine the possible effect of induced resistance (see experiment 3 below). Colour differences between LSR1 and P123/ LL01 allowed us to visually distinguish them in co-infestation experiments. The three clones were reared on *V. faba*, which is a favourable host for the various *A. pisum* biotypes (Boulain et al., 2019). Before each experiment, adult aphids were installed on *V. faba* for 24 hours to produce nymphs, and then removed. The nymphs were kept on the same plants until they reached the required age for the different experiments, and then transferred to pea for further analysis.

### Aphid performance assays

#### Experiment 1

We assumed that the effect of pea manipulation by the pea biotype may be limited to the vicinity of aphid feeding sites and may be transient. We first studied how the survival of LSR1 was affected by the pre-infestation of P123 in a clip cage (1.8 cm in diameter). Five P123 (five days old) aphids were confined in a clip cage on a fully expanded leaf of a two-week-old pea for three days and then replaced by six LSR1 (one day old) in the same cage. After seven days, the number of surviving LSR1 was counted. Pea plants with an empty clip cage were used as infestation control. Total 15 replicates were used for each treatment.

#### Experiment 2

We examined the effect of co-infestation by the pea biotype on the alfalfa biotype. In this experiment, three LSR1 (one day old) were co-infested with three P123 (one day old) in the same clip cage on a leaf of a two-week-old pea for seven days, and the surviving LSR1 were counted. This condition was compared with three LSR1 in a clip cage. Total 15 replicates were tested for each treatment.

#### Experiment 3

Because survival of LSR1 in the clip cages was low in the experiment 1 and 2, we examined the effect of P123 infestation on LSR1 performance on a whole pea plant where aphids could move freely. Ten LSR1 (one day old) mono-infested or co-infested with 10 P123, 20 P123 or 20 LL01 (one day old) were installed for seven days on a 10-day-old pea covered with an air-permeable plastic bag, and surviving aphids were then removed and counted. Three medium-sized LSR1 were selected from them and reinstalled on the same plant. The nymphs produced by the three LSR1 were then counted after two weeks. During the co-infestation experiment, the initial P123 or LL01 aphids were replaced by new one-day-old P123 or LL01 every week to avoid production of new larva and to maintain similar aphid density. Treatments with 10 or 20 P123 were used to analyse the effect of aphid density on LSR1 performance. Treatment with 20 LL01 was used to create the same aphid density effect as 20 P123 treatment without the expected effect of pea plant manipulation induced by the pea biotype P123. Total 30 replicates were examined for each treatment.

### Aphid feeding behaviour assays

Aphid feeding behaviour was monitored by the EPG technique (Tjallingii, 1978). A total of 20 aphids (one day old) composed of 20 P123 (control, pea-adapted), 20 LSR1 (control, pea-non-adapted) or 10 P123 and 10 LSR1 (co-infestation treatment) were installed on 10-day-old peas. After seven days of co-infestation, medium-sized aphids were selected from each of the three conditions and connected to a two cm long gold-wire with a small drop of conductive silver-glue (EPG Systems, Wageningen, Netherlands) on the dorsum and installed on the original pea plant on which the other aphids continued to feed. A reference electrode was inserted into soil. The electrical signals of aphid feeding behaviour were recorded by a direct current-EPG device (GIGA-8, EPG Systems, Wageningen, Netherlands). Each aphid was installed on the centre of the plant top and recorded for eight hours in a Faraday cage. The signals were analysed by Stylet^+^ software (EPG Systems, Wageningen, Netherlands). Only the data from aphids that spent more than 6 hours in the probing stage were analysed, yielding a total of 28 LSR1, 31 LSR1 and 26 P123 in the mono-infested LSR1, co-infestation and mono-infested P123 treatments, respectively.

Waveforms annotated in this study included non-probing (NP), probing (Pr), stylet pathway (C), intracellular punctures (pd), repetitive sieve element puncture (rpd), phloem salivation (E1), phloem sap ingestion (E2), derailed stylet (F) and xylem phase (G). Other phloem-associated parameters including fraction salivation (frE1, E1 followed by E2), single salivation (sgE1, E1 not followed by E2) and sustained ingestion (sE2, more than 10 min ingestion) were also calculated. The total number and duration of these waveforms were analysed by EPG-Calc (Giordanengo, 2014).

### Data analysis

All statistical analyses were conducted in R version 4.1.2 (R Core Team, 2021). The aphid performance traits were analysed against the experimental treatment (i.e. a fixed factor) by using Generalized linear mixed model with binomial and Poisson distribution for the survival rates and fecundity counts, respectively. As some experimental aphids were originated from the same replicate, the replicated ID was introduced as random factor to include this data dependency. Significance of the experimental treatment was estimated by using a likelihood ratio test. In case of significance, we analysed the estimated coefficients to identify factors levels that differed or not.

The number and total duration of each annotated EPG waveform were analysed against the experimental treatment (i.e. three levels fixed factors) by using Generalized linear models. While we used a Gamma model family to analyse the total duration of each EPG waveform, a Gaussian, Poisson or a Negative Binomial model family was used for analysing the number of the stages depending on data distribution and dispersion (Table S1). Significance of the experimental treatment was estimated by using a likelihood ratio test. As we performed multiple tests (one test per feeding behaviour), the p-value was corrected by using Benjamini Hocherg False Discovery Rate correction. In case of significance, the estimated coefficients were analysed to determine which experimental treatments differed. Transitional probabilities of feeding behaviours were analysed by using discrete time Markov chains modelling (markovchain R package) (Ebert et al., 2018; Spedicato, 2017). The median transition probability of each feeding stage were determined by taking the median value across all individuals in each treatment. An inspection of individual aphid’s feeding behaviour was made by a principal component analysis (PCA) with all the annotated waveforms except G phase, which was observed in only two aphids. Since we observed that co-infested LSR1 aphids split in two main groups on the PCA axes, we compared their EPG patterns using the same statistical analysis as presented above.

## Results

### Co-infestation but not pre-infestation induced susceptibility

We first confined aphids in clip cages and evaluated the effect of pre- and co-infestation of the pea biotype (clone P123) on the survival rate of the alfalfa biotype (clone LSR1). LSR1 survival rates were around 10% and were not significantly different on the leaves with or without a P123 pre-infestation of three days (Fig. 1A). In contrast, LSR1 co-infested with P123 showed a higher survival rate (around 50%) compared to mono-infested LSR1 (17.8%, Fig. 1B), which indicated that co-infestation with P123 in the same cage promoted LSR1 survival.

**Figure 1:**
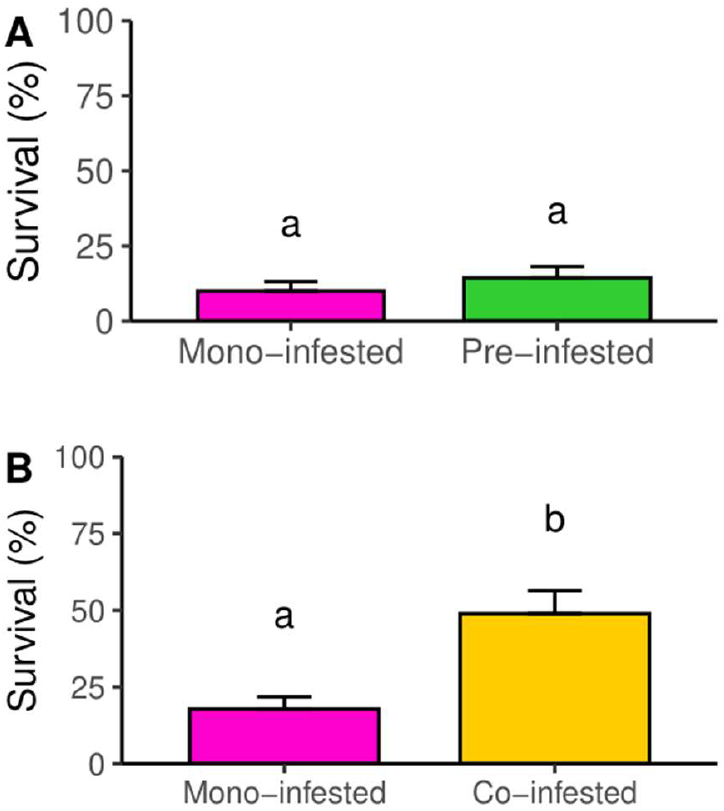
Influence of pre-infestation (A) or co-infestation (B) with the pea biotype (P123 clone) on survival rates of the alfalfa biotype (LSR1 clone) in a cage confinement. Results are presented as the mean ± standard error (n=15), and analysed by Generalized linear mixed model with binomial distribution. Different lower-case letters indicate significant difference between the treatments (P < 0.05).

As the aphids did not live long enough to reproduce in clip cages, we then examined LSR1 performance with P123 co-infestation on whole pea plants. LL01 (alfalfa biotype) was also co-infested to examine the effect of aphid density on the LSR1 performance. On the seventh day of co-infestation, the LSR1 survival rates were higher than 75% in all the treatments, and the LSR1 survival rates in both 20 LL01 and 10 P123 treatments were slightly higher than mono-infested LSR1 group (Fig. S1). Three LSR1 were kept for two additional weeks on the same pea. After 14 days from the initial installation, more than 50% of LSR1 survived on the P123-co-infested pea, while less than 25% of LSR1 survived in the LSR1 mono-infestation and the 20 LL01 co-infestation treatments (Fig. 2A). Three weeks after infestation, LSR1 adults produced around 20 nymphs on the P123-co-infested pea, but less than five nymphs when LSR1 were mono-infested or co-infested with 20 LL01 (Fig. 2B). No significant difference was observed between the 10 and the 20 P123 treatments. LSR1 with 20 LL01 showed a slightly better survival rate on day 7 than LSR1 mono-infestation, but this effect disappeared on day 14 and was not found for fecundity. Altogether, these results indicated that co-infestation with P123 increased the survival and fecundity of LSR1, while the co-infestation with LL01 had no effect on LSR1 survival, and that these effects depended little or not on aphid density.

**Figure 2:**
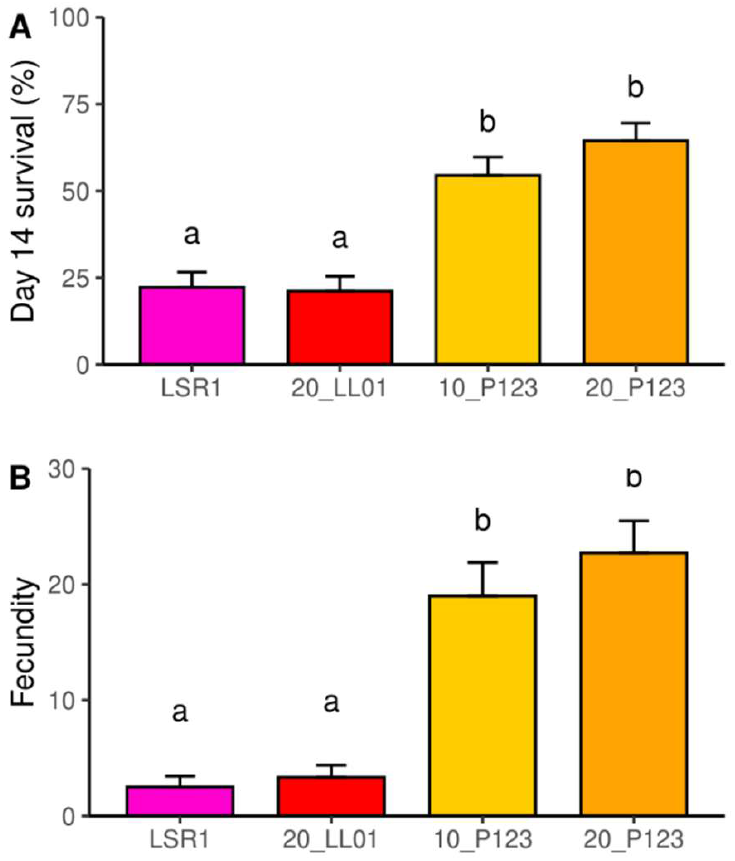
Influence of co-infestation with the pea biotype (P123 clone) or the alfalfa biotype (LL01 clone) on survival rates at day 14 post-infestation (A) and on cumulated fecundity at day 21 (B) of the alfalfa biotype (LSR1 clone) on whole pea plants. Results are presented as the mean ± standard error (n=30), and analysed by Generalized linear mixed model with binomial and Poisson distribution for the survival rates and fecundity, respectively. Different lower-case letters indicate significant differences between the treatments (P < 0.05).

### Co-infestation with the pea biotype increased phloem feeding time of the alfalfa biotype on pea

In order to understand how P123 promotes LSR1 fitness on pea, LSR1 feeding behaviours on pea were analysed by EPG with or without P123 co-infestation. While preparing the experiment, we confirmed that the LSR1 survival rate was significantly increased in the presence of P123, and that the P123 survival rate was not affected by LSR1 co-infestation (Fig. S2). The number and total duration of each annotated feeding stage of the EPG results are shown in Table 1. Because only two aphids showed xylem phase (G) (Table 1), G phase was excluded from further analysis.

**Table 1:**
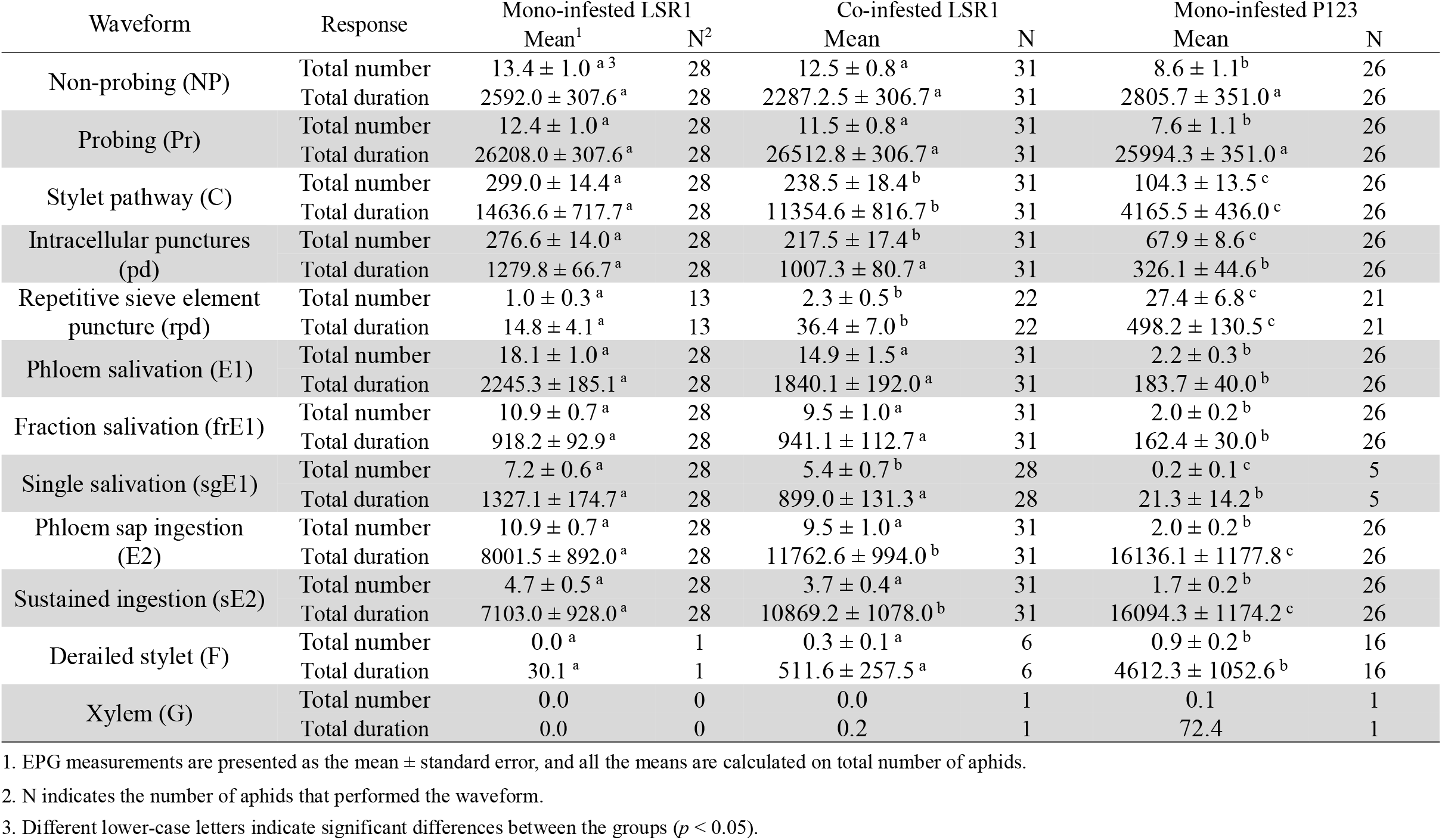
EPG variables measured on mono-infested LSR1 (n = 28), LSR1 co-infested with P123 (n = 31) and mono-infested P123 (n = 26).

| Waveform | Response | Mono-infested LSR1 |  | Co-infested LSR1 |  | Mono-infested P123 |  |
| --- | --- | --- | --- | --- | --- | --- | --- |
|  |  | Mean<sup>1</sup> | N<sup>2</sup> | Mean | N | Mean | N |
| Non-probing (NP) | Total number | 13.4 ± 1.0<sup>a3</sup> | 28 | 12.5 ± 0.8<sup>a</sup> | 31 | 8.6 ± 1.1<sup>b</sup> | 26 |
|  | Total duration | 2592.0 ± 307.6<sup>a</sup> | 28 | 2287.2.5 ± 306.7<sup>a</sup> | 31 | 2805.7 ± 351.0<sup>a</sup> | 26 |
| Probing (Pr) | Total number | 12.4 ± 1.0<sup>a</sup> | 28 | 11.5 ± 0.8<sup>a</sup> | 31 | 7.6 ± 1.1<sup>b</sup> | 26 |
|  | Total duration | 26208.0 ± 307.6<sup>a</sup> | 28 | 26512.8 ± 306.7<sup>a</sup> | 31 | 25994.3 ± 351.0<sup>a</sup> | 26 |
| Stylet pathway (C) | Total number | 299.0 ± 14.4<sup>a</sup> | 28 | 238.5 ± 18.4<sup>b</sup> | 31 | 104.3 ± 13.5<sup>c</sup> | 26 |
|  | Total duration | 14636.6 ± 717.7<sup>a</sup> | 28 | 11354.6 ± 816.7<sup>b</sup> | 31 | 4165.5 ± 436.0<sup>c</sup> | 26 |
| Intracellular punctures (pd) | Total number | 276.6 ± 14.0<sup>a</sup> | 28 | 217.5 ± 17.4<sup>b</sup> | 31 | 67.9 ± 8.6<sup>c</sup> | 26 |
|  | Total duration | 1279.8 ± 66.7<sup>a</sup> | 28 | 1007.3 ± 80.7<sup>a</sup> | 31 | 326.1 ± 44.6<sup>b</sup> | 26 |
| Repetitive sieve element puncture (rpd) | Total number | 1.0 ± 0.3<sup>a</sup> | 13 | 2.3 ± 0.5<sup>b</sup> | 22 | 27.4 ± 6.8<sup>c</sup> | 21 |
|  | Total duration | 14.8 ± 4.1<sup>a</sup> | 13 | 36.4 ± 7.0<sup>b</sup> | 22 | 498.2 ± 130.5<sup>c</sup> | 21 |
| Phloem salivation (E1) | Total number | 18.1 ± 1.0<sup>a</sup> | 28 | 14.9 ± 1.5<sup>a</sup> | 31 | 2.2 ± 0.3<sup>b</sup> | 26 |
|  | Total duration | 2245.3 ± 185.1<sup>a</sup> | 28 | 1840.1 ± 192.0<sup>a</sup> | 31 | 183.7 ± 40.0<sup>b</sup> | 26 |
| Fraction salivation (frE1) | Total number | 10.9 ± 0.7<sup>a</sup> | 28 | 9.5 ± 1.0<sup>a</sup> | 31 | 2.0 ± 0.2<sup>b</sup> | 26 |
|  | Total duration | 918.2 ± 92.9<sup>a</sup> | 28 | 941.1 ± 112.7<sup>a</sup> | 31 | 162.4 ± 30.0<sup>b</sup> | 26 |
| Single salivation (sgE1) | Total number | 7.2 ± 0.6<sup>a</sup> | 28 | 5.4 ± 0.7<sup>b</sup> | 28 | 0.2 ± 0.1<sup>c</sup> | 5 |
|  | Total duration | 1327.1 ± 174.7<sup>a</sup> | 28 | 899.0 ± 131.3<sup>a</sup> | 28 | 21.3 ± 14.2<sup>b</sup> | 5 |
| Phloem sap ingestion (E2) | Total number | 10.9 ± 0.7<sup>a</sup> | 28 | 9.5 ± 1.0<sup>a</sup> | 31 | 2.0 ± 0.2<sup>b</sup> | 26 |
|  | Total duration | 8001.5 ± 892.0<sup>a</sup> | 28 | 11762.6 ± 994.0<sup>b</sup> | 31 | 16136.1 ± 1177.8<sup>c</sup> | 26 |
| Sustained ingestion (sE2) | Total number | 4.7 ± 0.5<sup>a</sup> | 28 | 3.7 ± 0.4<sup>a</sup> | 31 | 1.7 ± 0.2<sup>b</sup> | 26 |
|  | Total duration | 7103.0 ± 928.0<sup>a</sup> | 28 | 10869.2 ± 1078.0<sup>b</sup> | 31 | 16094.3 ± 1174.2<sup>c</sup> | 26 |
| Derailed stylet (F) | Total number | 0.0<sup>a</sup> | 1 | 0.3 ± 0.1<sup>a</sup> | 6 | 0.9 ± 0.2<sup>b</sup> | 16 |
|  | Total duration | 30.1<sup>a</sup> | 1 | 511.6 ± 257.5<sup>a</sup> | 6 | 4612.3 ± 1052.6<sup>b</sup> | 16 |
| Xylem (G) | Total number | 0.0 | 0 | 0.0 | 1 | 0.1 | 1 |
|  | Total duration | 0.0 | 0 | 0.2 | 1 | 72.4 | 1 |

1. EPG measurements are presented as the mean ± standard error, and all the means are calculated on total number of aphids.
2. N indicates the number of aphids that performed the waveform.
3. Different lower-case letters indicate significant differences between the groups ( $p < 0.05$ ).

There was no significant difference in the total probing and non-probing time between all the three groups (Table 1 and Fig. S3B and D). However, P123 and LSR1 clones showed contrasting feeding behaviours in most other variables (Table 1 and Fig. S3). P123 probed less frequently than LSR1 did (Table1, Fig. S3A and C), indicating that P123 had longer period of probing and non-probing time compared to LSR1. P123 showed lower number of “stylet pathway” (C) and “intracellular punctures” (pd) phases compared with LSR1 (Table 1, Fig. S3E and G), and spent around 16% of the total recording time in the two phases (Fig. 3C). In comparison, LSR1 spent more than a half of recording time in C and pd phases before reaching phloem (Fig. 3A). Repetitive sieve element puncture (rpd) is reported to occur before the phloem phase starts and is more frequent in adapted biotype on its host plant than in non-adapted biotype (Schwarzkopf et al., 2013). Eighty one percent of P123 showed rpd, but only 46% of LSR1 did (Table 1). P123 also showed more frequent and longer rpd than LSR1 (Table 1, Fig. S3I and J). Aphids usually initiate phloem phase by a salivation phase (E1) before the ingestion phase (E2). The E1 phase can be then categorised into single E1 (sgE1), which is not followed by E2, and fractionated E1 (frE1), which is followed. Number and total duration of sgE1 and frE1 tend to increase in aphids feeding on resistant host plants (Tjallingii, 2006). Indeed, frE1 and sgE1 were more frequent and longer in LSR1 than in mono-infested P123 (Table 1). Once P123 started E1, there was a high transition rate (0.95) from E1 to E2, followed by a prolonged E2 phase (Fig. 3C). The total duration of the E2 phase in P123 averaged 16100 seconds and virtually all of these phloem sap ingestion phases were “sustained E2” (sE2) (Table 1), meaning that P123 can effectively establish a phloem feeding on pea. On the contrary, all LSR1 exhibited sgE1, and it accounted for more than 50% of the duration of E1 (Table 1). This caused lower transition rates from E1 to E2 (0.59) in LSR1 than in P123 (Fig. 3A). This, combined with a high transition probability (0.81) from E2 to E1 (Fig. 3A), resulted in LSR1 spending only 8001 seconds in E2 (Table 1). “Derailed stylet” (F) phase is usually interpreted as the result of stylet penetration difficulties, which are generally observed when aphids feed on non-adapted or resistant host plants (Denoirjean et al., 2021; González et al., 2022; Pompon & Pelletier, 2012). Surprisingly, more P123 (16 of total 26) showed F phase than LSR1 (one of total 28) (Table 1), and P123 also showed longer duration of F than LSR1 (Table1 and Fig. S3V).

**Figure 3:**
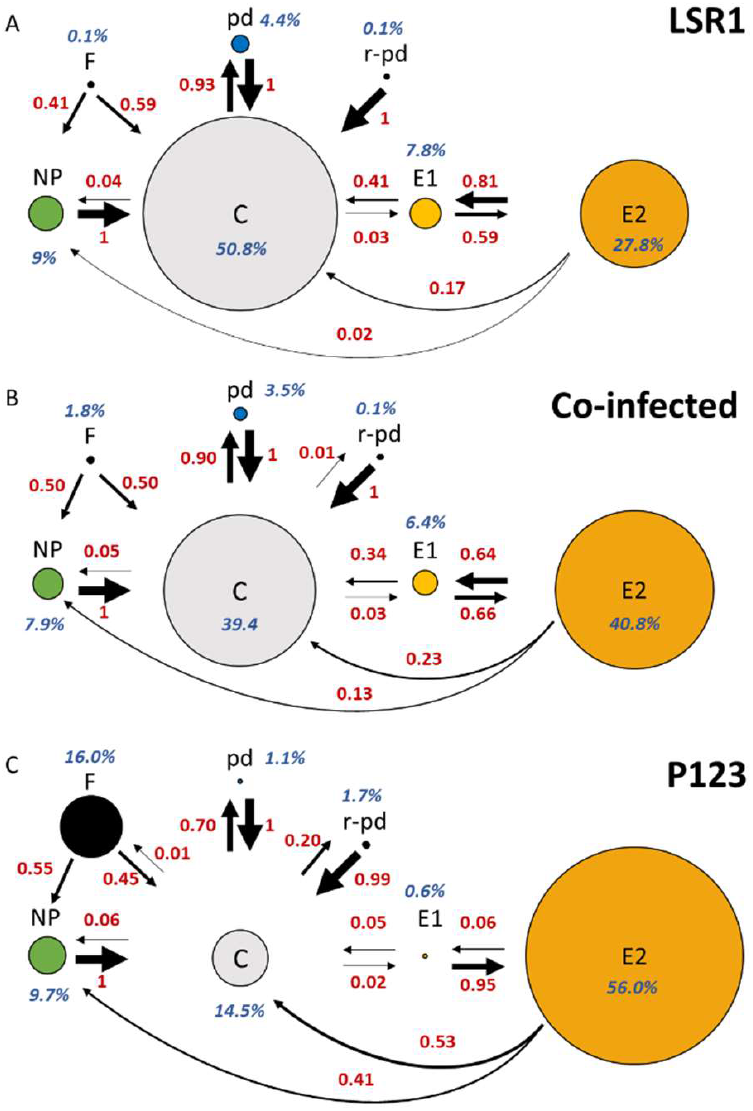
Transition probabilities of aphid feeding behaviours on pea in LSR1 in mono-infestation (A), LSR1 co-infested with P123 (B) and P123 in mono-infestation (C). The size of the circles is proportional to the time aphids spent in each recorded feeding stage that included non-probing (NP), stylet pathway (C), intracellular punctures (pd), repetitive sieve element puncture (rpd), phloem salivation (E1), phloem sap ingestion (E2) and derailed stylet (F). The thickness and value of arrows indicate the proportions of the behaviour transition. Arrows are not shown if the transition rates are lower than 0.01.

Compared with the feeding behaviour of mono-infested P123 or LSR1, LSR1 co-infested with P123 showed an intermediate pattern in many feeding phases. The co-infested LSR1 significantly reduced the number (Table 1, Fig. S3E) and total duration of C (around 40%, Table 1, Fig. 3B and Fig. S3F) and showed less pd (Table 1, Fig. S3G) compared with mono-infested LSR1. 71% of the co-infested LSR1 displayed rpd phase, which resulted in more number and longer total duration of rpd compared with the LSR1 mono-infestation (Table 1 and Fig. S3I and J). In addition, 90% of the co-infested LSR1 showed sgE1 compared to 100% in the mono-infested LSR1 group (Table 1). This indicated that co-infested LSR1 spent less time to reach phloem and had higher chance to reach E2 after E1 compared with the mono-infested LSR1. The transition from E2 to E1 was lower in the co-infestation group (0.64) than in the LSR1 group (0.81) (Fig. 3A and B), which resulted in a significantly longer duration of E2 in the co-infested LSR1 than in the mono-infested one (Fig. S3R and T). Interestingly, more LSR1 showed F phase when co-infested with P123 (Table 1). These results clearly indicated that LSR1 feeding behaviours were altered by P123 co-infestation and shifted toward those observed for the pea-adapted biotype.

The first component of the PCA, which accounts for 51.5% of total inertia, clearly separated P123 and LSR1 aphids under mono-infestation (Fig. 4 and S4). The projections of the feeding behaviours of the different LSR1 individuals in co-infestation with P123 were scattered among the ellipses formed by P123 and LSR1 mono-infestation groups, indicating that some LSR1 in co-infestation behaved similarly to LSR1 in mono-infestation, while others behaved more like P123. Based on the first and third axes of the PCA, we could assign seven LSR1 aphids of the co-infestation treatment to the group of P123 mono-infestation, while 24 were outside of the ellipse (Fig. S4A). Comparison between these two groups of co-infested LSR1 showed significant differences for most EPG variables except rpd and total durations of NP and Pr (Table S2), indicating that the feeding improvement by induced susceptibility was limited to some LSR1 aphids (23%). In particular, these co-infested LSR1 behaving like P123 showed longer duration of E2 and sE2 than the others (Table S2), which explains the extended E2 phase in the co-infestation group (Table 1 and Fig. S3R and T).

**Figure 4:**
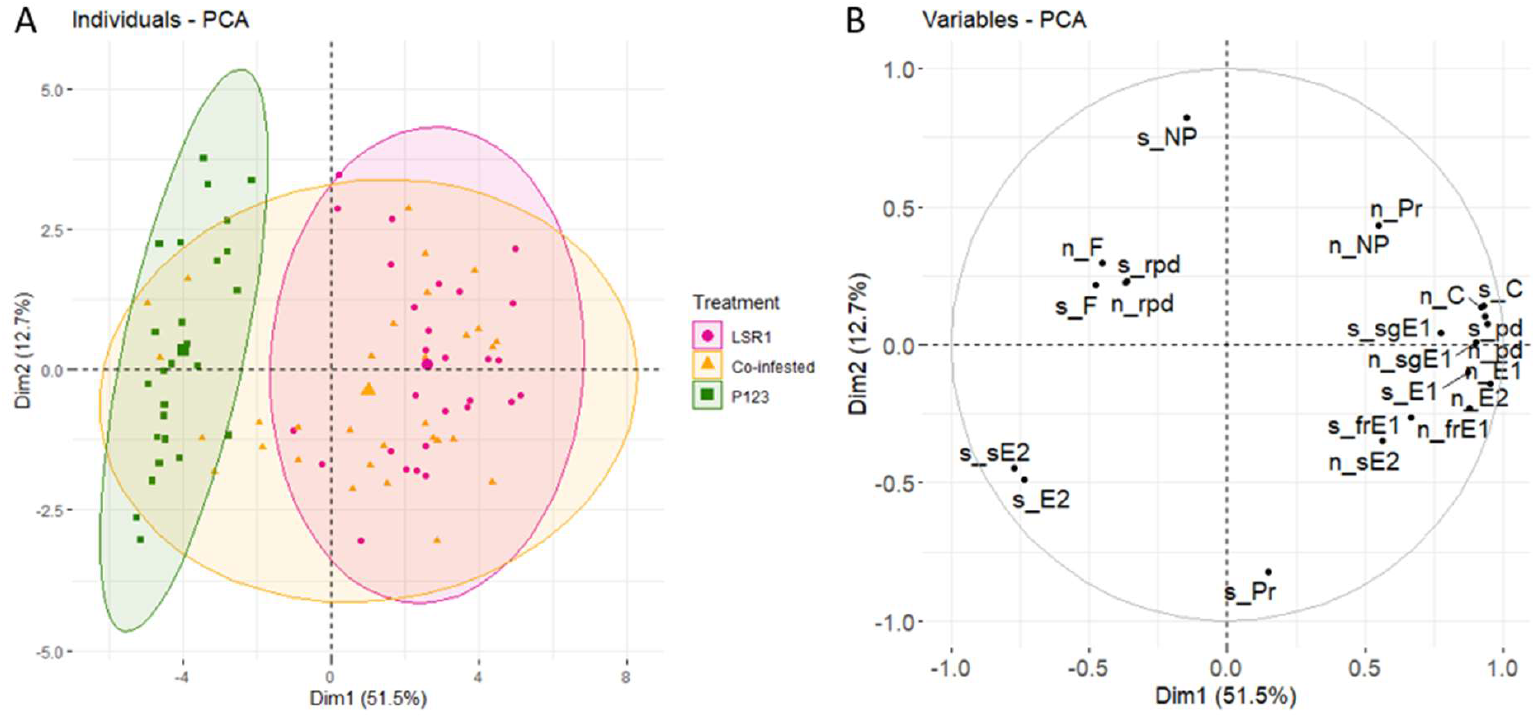
Principal component analysis (PCA) on selected EPG variables recorded for mono-infested LSR1 (pink spots, n=28), LSR1 co-infested with P123 (yellow spots, n=31) and mono-infested P123 (green spots, n=26). The circles show the confidence ellipses (with a confidence level of 0.95) calculated with the barycentre of individuals for each variable parameter on the factorial plan. (A): PCA of recorded individuals and (B): Variables factor map indicates the spatial ordination in dimensions 1 and 2. Selected variables are the number (n_) and the total duration (s_) of non-probing (NP), probing (Pr), stylet pathway (C), intracellular punctures (pd), phloem salivation (E1), fraction salivation (frE1), single salivation (sgE1), phloem sap ingestion (E2), and sustained ingestion (sE2) and derailed stylet (F).

## Discussion

In this study, we tested the hypothesis that *A. pisum* pea biotype manipulates pea plants and induces the susceptibility, allowing a non-adapted biotype to establish feeding and improve its performance on pea. We assumed that the effect of manipulation by the pea biotype was limited to the proximity of the feeding site and used clip cages to restrict the movement of the aphids. The survival rate of LSR1 was significantly increased in the P123 co-infestation condition but not in the P123 pre-infestation treatment. The results of co-infestation experiment indicated that P123 can induce pea susceptibility to LSR1 and promote its survival. The lack of observable effect of P123 pre-infestation may be due to the reversible effect of the P123-induced manipulation, probably due to the reactivation of plant defences. It could be also that three days of pre-infestation were not enough to induce sufficient pea susceptibility to LSR1. In the studies of *S. graminum* and *M. persicae*, the pre-infestation of virulent aphids were shown to increase the performance of avirulent aphids (Dorschner et al., 1987; Sun et al., 2020). Compared to these two examples, the manipulation caused by *A. pisum* pea biotype may be weaker and short-lasting, which could be explained by different mechanisms of manipulation. In the second experiment, the aphids could move freely on a whole plant, and LSR1 and P123 were not necessarily close to each other. Nevertheless, co-infestation with P123 showed significantly positive effect on the 14^th^ day survival rate and the fecundity of LSR1. Total fecundity of LSR1 in both 10 P123 and 20 P123 conditions did not show significant difference, which indicated that there might be a limitation of the P123-induced susceptibility to LSR1. Similar results were observed in a study of *A. glycines*-soybean interactions, which showed that co-infestation with two different densities of avirulent aphids increased the fecundity of both virulent and avirulent aphids without detecting a density effect (Varenhorst et al., 2015).

The co-infestation with 20 LL01 increased the survival rate of LSR1 at day seven, but the effect was no longer observed after the day 14^th^. Hence, the LSR1 fecundity increase caused by the P123 co-infestation was not due to the aphid density but due to the P123 specific effect on the pea plants. In the condition of 20 LL01, there were 30 alfalfa biotype aphids installed on a pea plant, but no negative effect on the LSR1 performance was observed compared to the LSR1 in mono-infestation. As in the comparison of 10 and 20 P123, increased number of the alfalfa biotype might not have additional negative effect on the LSR1 fecundity. Alternatively, we can also hypothesize that unlike some other examples (Ten Broeke et al., 2017; Züst & Agrawal, 2016), the *A. pisum* alfalfa biotype may trigger very little or no defence response in pea. This hypothesis is supported by the observation that co-infestation of equal number of LSR1 and P123 did not affect the survival of P123 (Fig. S2), although we can also reason that the manipulation by P123 is strong enough to overcome the pea defence triggered by the equal number of LSR1 aphids.

The EPG analyses of P123, LSR1 and P123-co-infested LSR1 aphids on pea plants revealed the behavioural changes of LSR1 feeding induced by the co-infestation. Analyses of EPG parameters showed that all the three groups showed similar total duration of probing but were different in many feeding stages. The differences between P123 and LSR1 mono-infestation groups were clearly distinguished by the navigation of stylets within pea plant, suggesting that pea deterrence to LSR1 exists in pea mesophyll and phloem cells and not in the plant surface, in accordance with earlier results (Schwarzkopf et al., 2013). The co-infestation with P123 decreased the duration of pathway phase, the numbers of intracellular punctures and salivation, and most importantly, increased the duration of phloem sap ingestion of LSR1 aphids. We can reasonably assume that the higher survival and fecundity the co-infested LSR1 aphids is a consequence of prolonged phloem feeding.

PCA analysis showed that the behaviours of co-infested LSR1 aphids were separated into two main groups, one resembling mono-infested P123 aphids and the other resembling mono-infested LSR1. In this EPG experiment, some LSR1 aphids could have been very close to P123 aphids and others not. The variable behaviour of co-infested LSR1 may reflect the variable distance between the LSR1 and surrounding P123 aphids. Alternatively, different plant parts may show specific responses to aphid infestation, and the freely moving LSR1 could feed on a susceptible part manipulated by P123. Aphid infestation on different plant parts can result in site-specific changes of phloem metabolites such as sugars and amino acids, which affect aphid performance differently (Jakobs et al., 2019).

The mechanisms by which pea-non-adapted LSR1 increased phloem feeding duration are unclear. In the case of pepper-*M. persicae* interaction, it was suggested that the induced susceptibility resulted from the suppression of ROS by virulent aphids (Sun et al., 2020). For the wheat-*S. graminum* interaction, induction of senescence-like damage on wheat leaves and short-term release of free amino acid by virulent aphid were observed (Dorschner et al., 1987). We did not observe any damage on the pea plants infested by *A. pisum* biotypes. Considering that the LSR1 co-infested with P123 showed longer phloem feeding duration, it is likely that P123 changed accessibility or condition of phloem so that LSR1 can keep feeding, rather than they changed the nutritional quality of the pea phloem. A very short salivation of P123 could achieve a long phloem feeding that follows, while LSR1 showed repeated salivation and short phloem feeding phases. These behavioural differences indicated that P123 either injected a greater amount of effective salivary components (e.g. proteins, RNAs) than non-adapted biotypes in short periods of time or injected pea-biotype specific components to sustain the phloem feeding in pea. Indeed, pea and alfalfa biotypes differ in some of their salivary genes both in expression and sequence (Boulain et al., 2019; Nouhaud et al., 2014). Analyses of plant physiology and further functional characterization of salivary components may help to understand the mechanisms of induced susceptibility in pea by pea-adapted A. pisum. In this study, we observed an induced susceptibility to the alfalfa biotype, but we do not know if the pea biotype also induces pea susceptibility to other *A. pisum* biotypes or other aphid species that do not feed on pea. Further investigation of pea-biotype induced susceptibility may reveal different types of pea and aphid interactions and the pea defence mechanisms against other aphids.

## Acknowledgements

We would like to thank Marie-Laure Pilet-Nayel, Isabelle Glory and Angélique Lesné for providing the pea seeds.

