## Supplemental Figure 1 for "Pea-adapted biotype of the aphid *Acyrthosiphon pisum* induces susceptibility of pea to non-adapted biotype enabling improved feeding and performance"

**Supplemental data**

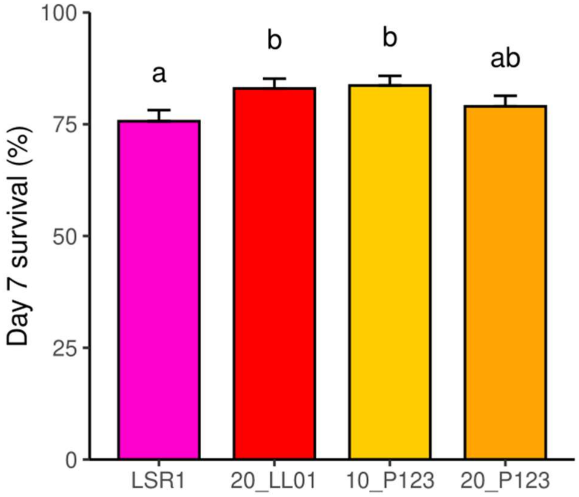


Figure S1: Influence of co-infestation with the pea biotype (P123 clone) or the alfalfa biotype (LL01 clone) on survival rates of the alfalfa biotype (LSR1 clone) on day 7 on whole pea plants. Results are presented as the mean ± standard error (n=30), and analysed by Generalized linear mixed model with binomial and Poisson distribution for the survival rates and fecundity, respectively. Different lower-case letters indicate significant differences between the treatments ($P < 0.05$).
