## Supplemental Figure 2 for "Pea-adapted biotype of the aphid *Acyrthosiphon pisum* induces susceptibility of pea to non-adapted biotype enabling improved feeding and performance"

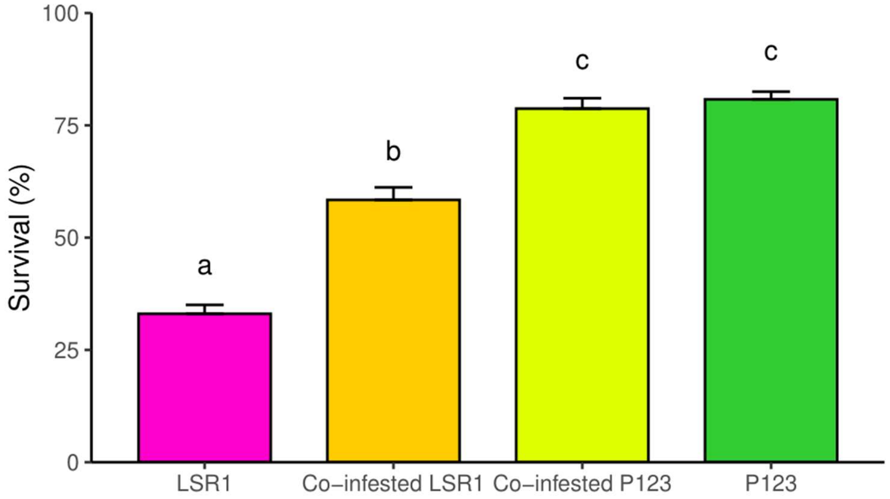


Figure S2: Influence of co-infestation with the pea biotype (P123 clone) on survival rates of the alfalfa biotype (LSR1 clone) on whole pea plants in the EPG experiment. The live LSR1 numbers in the LSR1 mono-infested (n=28) and co-infested (n=31) groups and the live P123 numbers in the P123 mono-infested (n=26) and co-infested (n=31) groups were counted at day 7 post-infestation. Results are presented as the mean ± standard error, and analysed by Generalized linear mixed model with binomial distribution. Different lower-case letters indicate significant difference between the treatments ($P < 0.05$).
