## Supplemental Figure 3 for "Pea-adapted biotype of the aphid *Acyrthosiphon pisum* induces susceptibility of pea to non-adapted biotype enabling improved feeding and performance"

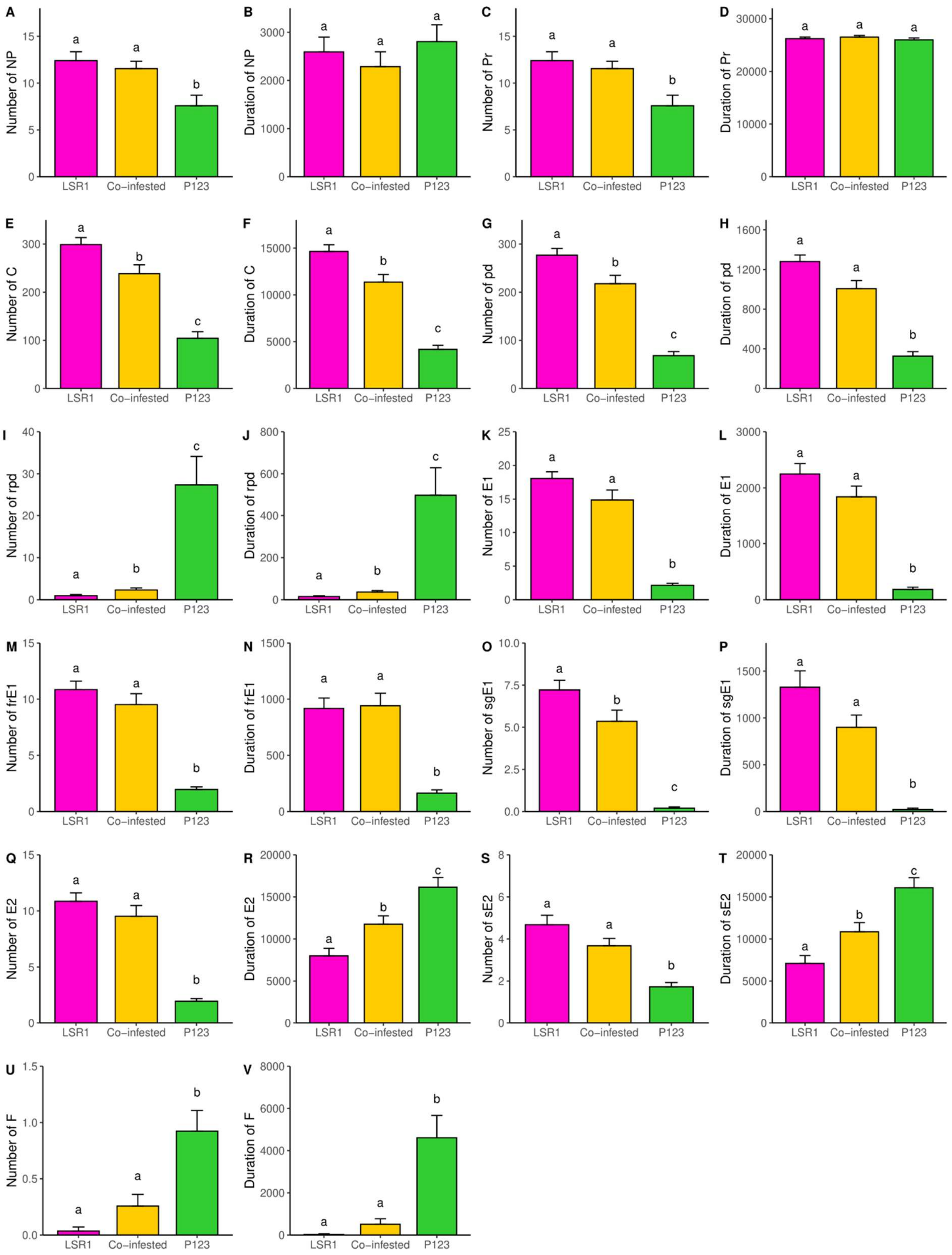


Figure S3: Number and total duration in second of variables of LSR1 mono-infestation, LSR1 co-infested with P123 and P123 mono-infestation measured by Electrical Penetration Graph.

EPG variables included non-probing (A and B), probing (C and D), stylet pathway (E and F), intracellular punctures (G and H), repetitive sieve element puncture (I and J), phloem salivation (K and L), fraction salivation (M and N), single salivation (O and P), phloem sap ingestion (Q and R), and sustained ingestion (S and T) and derailed stylet (U and V) and are presented as the mean ± standard error, and all the means are calculated on total number of aphids. Different lower-case letters indicate significant differences between the groups ($p < 0.05$).
