## Supplemental Figure 4 for "Pea-adapted biotype of the aphid *Acyrthosiphon pisum* induces susceptibility of pea to non-adapted biotype enabling improved feeding and performance"

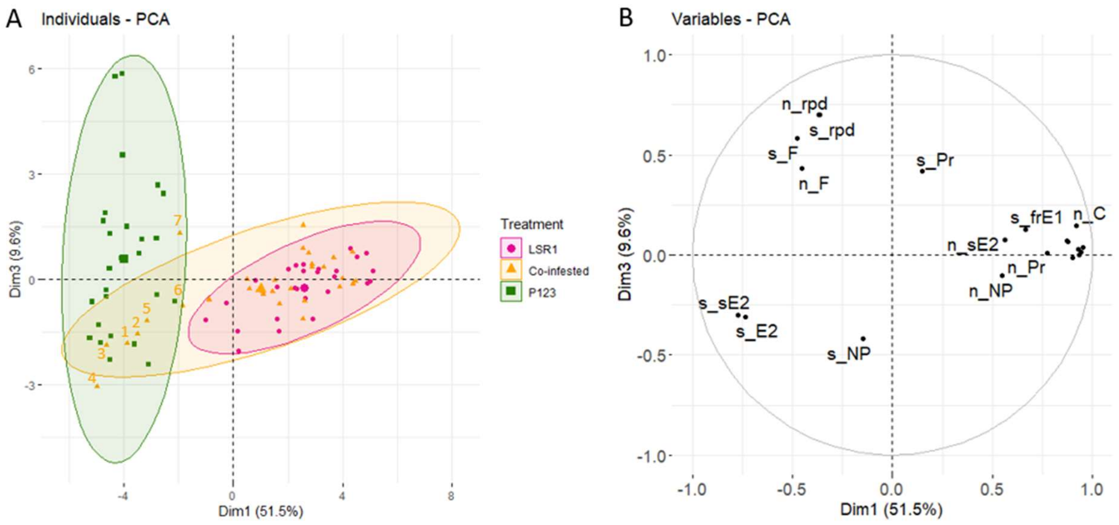


Figure S4: Principal component analysis (PCA) on selected EPG variables recorded for mono-infested LSR1 (pink spots, n=28), LSR1 co-infested with P123 (yellow spots, n=31) and mono-infested P123 (green spots, n=26). The circles show the confidence ellipses (with a confidence level of 0.95) calculated with the barycentre of individuals for each variable parameter on the factorial plan. (A): PCA of recorded individuals and (B): Variables factor map indicates the spatial ordination in dimensions 1 and 3. The seven co-infested LSR1 inside P123 ellipse are labelled, which include the five individuals shown in the P123 ellipse of Figure 4A. Selected variables are the number (n_) and the total duration (s_) of non-probing (NP), probing (Pr), stylet pathway (C), intracellular punctures (pd), phloem salivation (E1), fraction salivation (frE1), single salivation (sgE1), phloem sap ingestion (E2), and sustained ingestion (sE2) and derailed stylet (F).
