## Supplemental Table 1 for "Pea-adapted biotype of the aphid *Acyrthosiphon pisum* induces susceptibility of pea to non-adapted biotype enabling improved feeding and performance"

Table S1: Statistical models and results used for analysing EPG variables

| EPG Waveform | Response | Model used | Statistics | df | Corrected *p*-value |
|---|---|---|---|---|---|
| NP | Total number | NB Model | $\chi^2$=15.07 | 2 | 0.001 |
| | Total duration | Gamma Model | F=0.64 | 2 | 0.532 |
| Pr | Total number | NB Model | $\chi^2$=14.73 | 2 | 0.001 |
| | Total duration | Gamma Model | F=0.64 | 2 | 0.532 |
| C | Total number | Gaussian Model | F=36.98 | 2 | <0.001 |
| | Total duration | Gamma Model | F=41.70 | 2 | <0.001 |
| pd | Total number | Gaussian Model | F=53.43 | 2 | <0.001 |
| | Total duration | Gamma Model | F=40.27 | 2 | <0.001 |
| rpd | Total number | NB Model | $\chi^2$=82.68 | 2 | <0.001 |
| | Total duration | Gamma Model | F=40.63 | 2 | <0.001 |
| E1 | Total number | NB Model | $\chi^2$=174.93 | 2 | <0.001 |
| | Total duration | Gamma Model | F=56.27 | 2 | <0.001 |
| frE1 | Total number | NB Model | $\chi^2$=113.92 | 2 | <0.001 |
| | Total duration | Gamma Model | F=35.59 | 2 | <0.001 |
| sgE1 | Total number | NB Model | $\chi^2$=159.57 | 2 | <0.001 |
| | Total duration | Gamma Model | F=11.56 | 2 | <0.001 |
| E2 | Total number | NB Model | $\chi^2$=113.92 | 2 | <0.001 |
| | Total duration | Gamma Model | F=12.69 | 2 | <0.001 |
| sE2 | Total number | Poisson Model | $\chi^2$=38.75 | 2 | <0.001 |
| | Total duration | Gamma Model | F=12.19 | 2 | <0.001 |
| F | Total number | Poisson Model | $\chi^2$=30.27 | 2 | <0.001 |
| | Total duration | Gamma Model | F=6.57 | 2 | 0.002 |

NB: Negative Binomial.
