## Supplemental Table 2 for "Pea-adapted biotype of the aphid *Acyrthosiphon pisum* induces susceptibility of pea to non-adapted biotype enabling improved feeding and performance"

Table S2: Comparison of the feeding behaviours of co-infested LSR1 aphids in the ellipse formed by mono-infested P123 (Group 1, n=7) and the remaining co-infested LSR1 (Group 2, n=24).

| EPG Waveform | Response | Model used | Statistics | Corrected *p*-value | Comparison |
|---|---|---|---|---|---|
| NP | Total number | Poisson Model | $c^2$=9.79 | 0.003 | Group 2 > Group 1 |
| | Total duration | Gamma Model | F=2.18 | 0.167 | Group 2 = Group 1 |
| Pr | Total number | Poisson Model | $c^2$=10.72 | 0.002 | Group 2 > Group 1 |
| | Total duration | Gamma Model | F=2.37 | 0.158 | Group 2 = Group 1 |
| C | Total number | Gaussian Model | F=70.22 | <0.001 | Group 2 > Group 1 |
| | Total duration | Gamma Model | F=45.05 | <0.001 | Group 2 > Group 1 |
| pd | Total number | Gaussian Model | F=67.56 | <0.001 | Group 2 > Group 1 |
| | Total duration | Gamma Model | F=72.08 | <0.001 | Group 2 > Group 1 |
| rpd | Total number | Poisson Model | $c^2$=0.13 | 0.758 | Group 2 = Group 1 |
| | Total duration | Gamma Model | F=0.01 | 0.932 | Group 2 = Group 1 |
| E1 | Total number | Poisson Model | $c^2$=96.89 | <0.001 | Group 2 > Group 1 |
| | Total duration | Gamma Model | F=28.46 | <0.001 | Group 2 > Group 1 |
| frE1 | Total number | Poisson Model | $c^2$=58.41 | <0.001 | Group 2 > Group 1 |
| | Total duration | Gamma Model | F=10.22 | 0.005 | Group 2 > Group 1 |
| sgE1 | Total number | Poisson Model | $c^2$=52.42 | <0.001 | Group 2 > Group 1 |
| | Total duration | Gamma Model | F=10.14 | 0.005 | Group 2 > Group 1 |
| E2 | Total number | Poisson Model | $c^2$=48.68 | <0.001 | Group 2 > Group 1 |
| | Total duration | Gamma Model | F=17.46 | <0.001 | Group 1 > Group 2 |
| sE2 | Total number | Poisson Model | $c^2$=7.93 | 0.006 | Group 2 > Group 1 |
| | Total duration | Gamma Model | F=17.46 | <0.001 | Group 1 > Group 2 |
